# Evaluation of the functional role of the maize *Glossy2 and Glossy2-like* genes in cuticular lipid deposition

**DOI:** 10.1101/2020.02.27.968321

**Authors:** Liza Esther Alexander, Yozo Okazaki, Michael A. Schelling, Aeriel Davis, Xiaobin Zheng, Ludmila Rizhsky, Marna D. Yandeau-Nelson, Kazuki Saito, Basil J. Nikolau

**Affiliations:** Roy J. Carver Department of Biochemistry, Biophysics and Molecular Biology, Iowa State University, Ames, IA, USA; Center for Metabolic Biology, Iowa State University, Ames, IA, USA; RIKEN Center for Sustainable Resource Science, Tsurumi-ku, Yokohama, Kanagawa 230-0045, Japan; Departments of Genetics, Development, and Cell Biology, Iowa State University, Ames, IA, USA; Graduate School of Pharmaceutical Sciences, Chiba University, Chuo-ku, Chiba 260-8675, Japan

## Abstract

Plant epidermal cells express unique molecular machinery that juxtapose the assembly of intracellular lipid components and the unique extracellular cuticular lipids that are unidirectionally secreted to plant surfaces. In maize (*Zea mays* L.), mutations at the *glossy2 (gl2)* locus affect the deposition of extracellular cuticular lipids. Sequence-based genome scanning identified a novel *gl2* homolog in the maize genome, *Gl2-like*. Sequence homology identifies that both the *Gl2-like* and *Gl2* genes are members of the BAHD superfamily of acyltransferases, with close sequence homology to the Arabidopsis *CER2* gene. Transgenic experiments demonstrate that *Gl2-like* and *Gl2* functionally complement the Arabidopsis *cer2* mutation, with differential impacts on the cuticular lipids and the lipidome of the plant, particularly affecting the longer alkyl chain acyl lipids, particularly at the 32-carbon chain length. Site-directed mutagenesis of the putative BAHD catalytic HXXXDX-motif indicates that *Gl2-like* requires this catalytic capability to fully complement the *cer2* function, but *Gl2* can accomplish this without the need for this catalytic motif. These findings demonstrate that both *Gl2* and *Gl2-like* overlap in their cuticular lipid function, however the two genes have evolutionary diverged to acquire non-overlapping functions.

**One-sentence summary:** Transgenesis dissection of the functional roles of the maize *Glossy2* and *Glossy2-Like* genes in cuticular lipid deposition.

## INTRODUCTION

Extracellular lipids are constituents of a protective hydrophobic structure, the cuticle, which covers aerial organs of all terrestrial plants. The cuticle plays important roles in many plant-environment interactions, including controlling water status (Kerstiens, 1996; Riederer, 2006), responding to abiotic stresses (Long, 2003), interacting with biotic pathogens (Kolattukudy, 1985; Jenks et al., 1994), and in defining organ boundaries during development (Yephremov et al., 1999; Sieber et al., 2000; Kurdyukov et al., 2006). The extracellular cuticular lipids are chemically and physically arranged in distinct layers (Kolattukudy, 1965; Jellings and Leech, 1982; Jetter et al., 2000), with the epicuticular lipids being primarily linear alkyl chains, including very-long chain fatty acids (VLCFAs), hydrocarbons, ketones, primary and secondary alcohols, aldehydes and wax esters; in addition, non-alkyl, terpene-type specialized metabolites are also components of the epicuticle (Martin and Juniper, 1970; Kolattukudy, 1976; Tulloch 1976). These lipids are deposited on and embedded within a lipophilic cutin polymer matrix (a polymer of esterified ω−hydroxy and epoxy-C16 and C18 fatty acids, glycerol, and α,ω−dicarboxylic acids) (Kolattukudy, 2001; Heredia, 2003; Bonaventure et al., 2004; Franke et al., 2005; Pollard et al., 2008).

The molecular aspects of cuticular lipid biogenesis have been greatly facilitated by forward genetic approaches that use *eceriferum (cer)* mutants of Arabidopsis (Koornneef et al., 1989), *glossy (gl)* mutants of maize (Schnable et al., 1994) and tomato (Vogg et al., 2004; Leide et al., 2007) and *wax crystal-sparse leaf (wsl)* mutants of rice (Yu et al., 2008; Wang et al., 2017). Results of these studies have been primarily interpreted in the context of a metabolic model that had been proposed from earlier physiological/biochemical studies (Bianchi et al., 1985; Post-Beittenmiller, 1996). At the core of this metabolic model is the endoplasmic reticulum associated process of fatty acid elongation, which feeds two fatty acid modification pathways: a reductive pathway that generates fatty aldehydes, primary alcohols, and wax esters; and a decarbonylative pathway that converts the common aldehyde intermediate to hydrocarbons, and ultimately ketones, and secondary alcohols.

In maize both fatty acid modification pathways are differentially expressed among different organs. Specifically, in juvenile leaves of maize seedlings, the reductive pathway predominates, and the expression of this metabolic network is under the control of the juvenile-to-adult phase transition, mediated by the transcription factor *Gl15* (Moose and Sisco, 1996). In contrast, the fatty acid elongation-decarboxylative pathway is primarily expressed in silks (Perera et al., 2010; Loneman et al., 2017; Dennison et al., 2019). The *gl* mutations that have been characterized since the early-1900s have been identified via phenotypic screens of seedling leaves (Hayes and Brewbaker, 1928), which therefore primarily affect the fatty acid elongation-reductive pathway.

*Glossy2* (*ZmGl2*) is exemplary of such a product of forward genetics, initially identified in 1928 as a mutant that causes “beading” of water droplets on seedling leaves, a common phenotype of all *gl* mutants (Hayes and Brewbaker, 1928). The initial characterization of the cuticular lipid profiles of *gl2* (Bianchi, 1975) suggested deficiencies in lipids that are derived from the fatty acid elongation-reductive pathway, specifically a potential block in the fatty acid elongation process between chain lengths of 30 and 32 carbon atoms. The homology of *ZmGl2* to the Arabidopsis *CER2* gene became apparent when both loci were molecularly defined (Tacke et al., 1995; Xia et al., 1996); the two genes encode proteins that share ∼60% sequence similarity. The *cer2* mutation causes the bright-green appearance of Arabidopsis stems (Koornneef et al., 1989), due to an underlying decrease in the products of the fatty acid elongation-decarboxylative pathway, with an apparent block in the elongation of C26 or C28 fatty acids to C30 fatty acid (Mcnevin et al., 1991; Jenks et al., 1995).

At the time of initial identification, these two proteins (ZmGL2 and CER2) defined novel sequences (Tacke et al., 1995; Xia et al., 1996), which ultimately became archetypal of the BAHD class of enzymes (St-Pierre and Luca, 2000; D’Auria, 2006). BAHD enzymes are acyl-CoA-dependent acyltransferases that catalyze the acylation of alcohols or amine groups, forming ester and amide bonds in the assembly of a large number of specialized metabolites (St-Pierre and Luca, 2000; D’Auria, 2006). These enzymes have been phylogenetically categorized into five clades (Clades I-V), with ZmGL2 and CER2 being members of Clade II (D’Auria, 2006).

The specific biochemical functions of the GLOSSY2/CER2-containing Clade II BAHD enzymes remains unclear. More recently, yeast reconstitution experiments suggest that CER2 (Haslam et al., 2012) and its rice homolog, OsCER2 (Wang et al., 2017) associate with the fatty acid elongase (FAE) system, but the role of the BAHD acyltransferase catalytic capability of these proteins is unclear.

In this study, we expanded the *Gl2* characterization by identifying a novel *Glossy2-like* (*ZmGl2-like*) gene in the maize genome that is a homolog of *ZmGl2,* which suggests that these two genes are products of an evolutionary gene duplication event that may have provided a template for potential neofunctionalization (Renny-Byfield and Wendel, 2014). Specifically, we employed a transgenic strategy to evaluate and compare the functional ability of *ZmGl2* and *ZmGl2-like* to genetically complement the Arabidopsis *cer2* mutant. The resulting transgenic lines were used to evaluate the effects of these genetic manipulations on the cuticular lipid profiles and also on the broader lipid metabolic network (i.e., cutin and cellular lipidome). These experiments demonstrate that both the maize *ZmGl2* and *ZmGl2-like* genes can complement the Arabidopsis *cer2* mutant, however, the two genes have functionally drifted, with *ZmGl2* expressing additional capabilities as compared to *ZmGl2-like* and *CER2*.

## RESULTS

### Computational identification and characterization of the *ZmGl2-like* gene

The phylogenetic diversity of the *ZmGl2* (GRMZM2G098239; Zm00001d002353) gene was explored by sequence-based homology searches of both the Arabidopsis and maize genomes (Altschul et al., 1997; Alonso et al., 2003; Andorf et al., 2016). This revealed that *ZmGl2* shares high sequence homology with an uncharacterized maize gene, we term *ZmGl2-like* (GRMZM2G315767; Zm00001d024317); this gene also shares homology with three Arabidopsis genes, *CER2* (At4g24510), *CER2-LIKE1* (At4g13840), and *CER2-LIKE2* (At3g23840) (Figure 1; Table 1) (Xia et al., 1996; Pascal et al., 2013). The ZmGL2 and ZmGL2-LIKE proteins share 50% sequence identity with each other, and ∼30% sequence identities with the CER2-family of proteins. This sequence homology includes the conserved HXXXDX, acyl-CoA-dependent acyl transferase domain, which is typical of the BAHD enzyme family (D’Auria, 2006).

**Figure 1.**
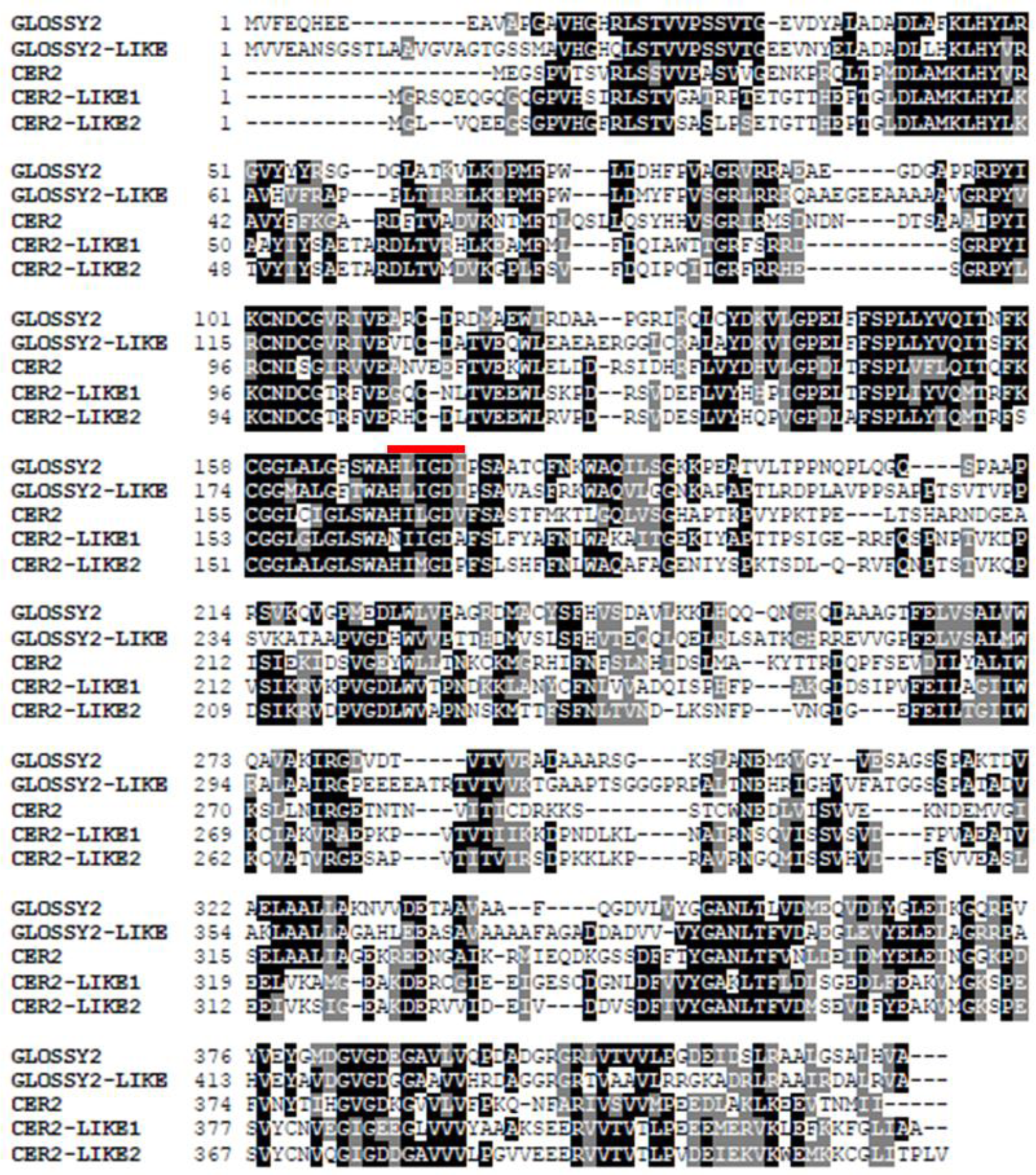
Amino acid sequence comparison of GLOSSY2 homologs. The maize GLOSSY2 (GRMZM2G098239; Zm00001d002353) and GLOSSY2-LIKE (GRMZM2G315767; Zm00001d024317) sequences are compared to the Arabidopsis CER2 (At4g24510), CER2-LIKE1 (At4g13840) and CER2-LIKE2 (At3g23840) sequences using Clustal O (1.2.4) and BOXSHADE (v3.21). Black shading identifies identical residues, and gray-shading identifies similar residues. The conserved -HXXXDX-acyl-CoA-dependent acyltransferase catalytic-domain of BAHD enzymes is identified with a red line.

**Table 1.**
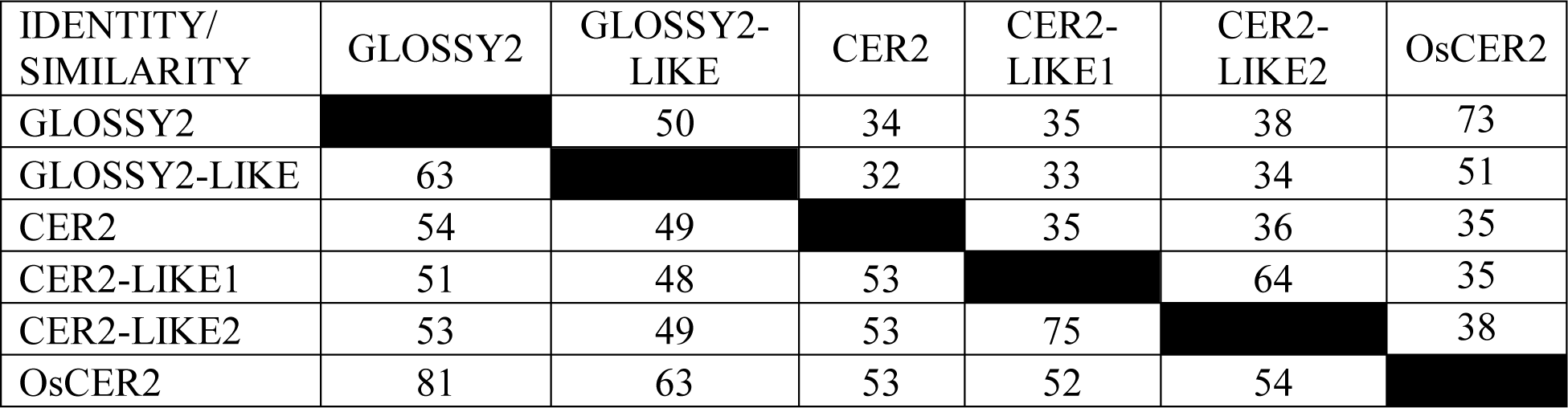
Conservation in the amino acid sequences of Zea mays GLOSSY2 homologs, CER2 homologs and rice OsCER2. The digits above the diagonal are % identity and the digits below the diagonal are % similarity.

### Transgenic expression of ZmGL2 and ZmGL2-LIKE modifies the *eceriferum* phenotype of the Arabidopsis *cer2* mutant stems

The functionality of the *ZmGl2* and *ZmGl2-like* genes was evaluated by their transgenic ability to complement the *cer2* mutant of Arabidopsis. These effects are interpreted in the context of the effect of removing such functionality in the homologous host, i.e., the maize *gl2* mutant (note a mutation at the *gl2-like* locus is not currently available). The *gl2* mutation primarily affects the accumulation of the major components of the cuticular lipids on maize seedling leaves, reducing the levels of C32 primary alcohol and C32 aldehyde, which are associated with only a partial compensatory increase in C28 primary alcohol and aldehyde (Supplemental Figure 1, Supplemental Table 1).

The ORFs coding for the ZmGl2 and ZmGl2-like proteins were expressed in Arabidopsis under the transcriptional control of the constitutive 35S promoter in homozygous lines that carried either wild-type or mutant *cer2-5* alleles. Expression of the *ZmGl2* and *ZmGl2-like* transgenes was confirmed at the protein and mRNA levels by western blot analysis (Figure 2A) of extracts using a GLOSSY2 antibody, and by RT-PCR analysis of RNA isolated from these plants (Figure 2B), respectively. As expected, the GL2 antibody detects a 46-kDa-polypeptide band in extracts from maize silk tissues, and this protein band is also detected in extracts prepared from Arabidopsis transgenic lines that are expressing the *Gl2* transgene (i.e., genotype: *ZmGl2* in WT and *ZmGl2* in *cer2-5*); this protein band is absent from the control, non-transgenic Arabidopsis plants. *ZmGl2-like* expression was detected via RT-PCR analysis with RNA-template preparations made from Arabidopsis plants carrying the *ZmGl2-like* transgene (in the WT and *cer2-5* mutant lines), and this transcript was undetectable in the non-transgenic wild-type and *cer2-5* mutant control plants (Figure 2B).

**Figure 2.**
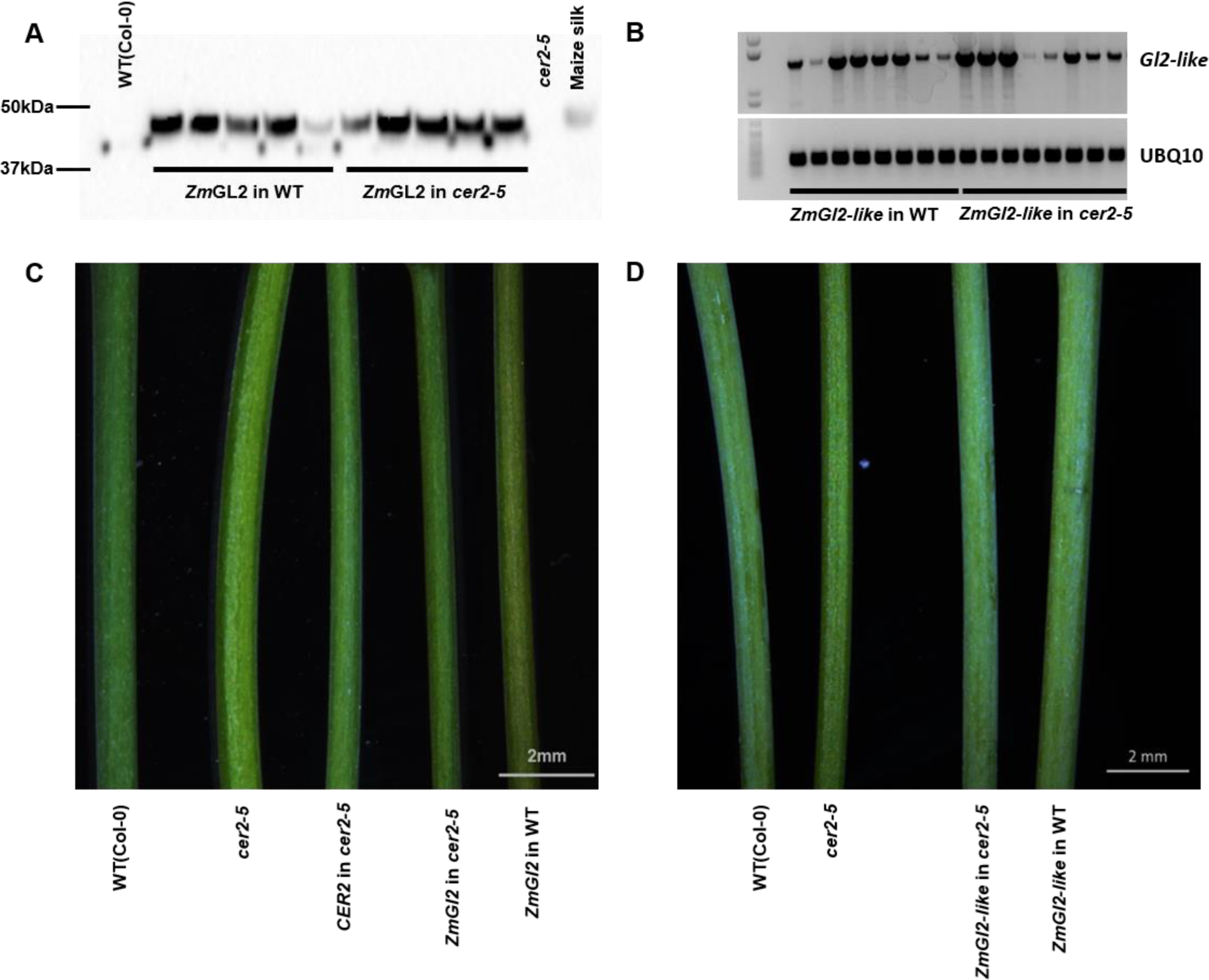
Transgenic expression of *ZmGl2* and *ZmGl2-like* in Arabidopsis. A) Western blot analysis of the 46-kDa GLOSSY2 protein transgenically expressed in either wild-type or cer2-5 mutant plants. Protein extract sample from maize silks serves as the positive control, protein extracts from the stems of wild-type Arabidopsis, and cer2-5 mutant plants serve as negative controls. B) RT-PCR analysis of the ZmGl2-like mRNA transgenically expressed in either wild-type or cer2-5 mutant plants. Ubiquitin mRNA (At4g05320) is used as the internal control. C) Stem phenotypes of non-transgenic Arabidopsis wild-type and cer2-5 mutant, and transgenic lines expressing either maize Gl2 or CER2 in wild-type and cer2-5 mutant backgrounds. D) Stem phenotypes of non-transgenic Arabidopsis wild-type and cer2-5 mutant, and transgenic lines expressing the maize Gl2-like transgene in either wild-type or cer2-5 mutant backgrounds.

Typical of *cer* mutants, the stems of the *cer2-5* mutant show the *eceriferum* phenotype, presenting bright green stems (Koornneef et al., 1989), rather than the dull green appearance of the wild-type plants (Figure 2C and 2D). This phenotype is associated with epicuticle-deficiency and is indicative of changes in the cuticular surface lipid composition (Koornneef et al., 1989). As with the transgenic expression of the CER2 protein (Xia et al., 1996), the transgenic expression of the ZmGL2-LIKE protein in the *cer2-5* mutant background fully restores the stem *eceriferum* phenotype to the dull green, wild-type appearance (Figure 2D). In contrast, the transgenic expression of the ZmGL2 protein in the *cer2-5* mutant only partially restored this phenotype to the wild-type appearance (Figure 2C). As control experiments, we transgenically expressed the *ZmGl2* or *ZmGl2-like* transgenes in wild-type Arabidopsis plants, and this did not alter the phenotype of the stems (i.e., they retained the dull green, wild-type appearance; Figure 2C and 2D).

As is expected from prior studies (Xia et al., 1996), scanning electron microscopic (SEM) examination of the stems identify the crystalloid structures of the epicuticle, and these are absent from the surfaces of Arabidopsis *cer2-*5 mutant stems, but the transgenic expression of *CER2* in this mutant background induces their reappearance and they have similar structures to those of wild-type plants (Figure 3). The parallel SEM examination of the stem-surfaces of plants expressing the *ZmGl2-like* transgene in either the wild-type or *cer2-5* mutant background indicate that the epicuticle is similar to that of wild-type plants (Figure 3). In contrast, the crystalloids on those plants expressing the *ZmGl2* transgene, in either the wild-type or *cer2-5* mutant background, occur at a lower density, and the structures of these crystalloids are more irregular, flattened, and have a thinner flake-like appearance than the wild-type control (Figure 3).

**Figure 3.**
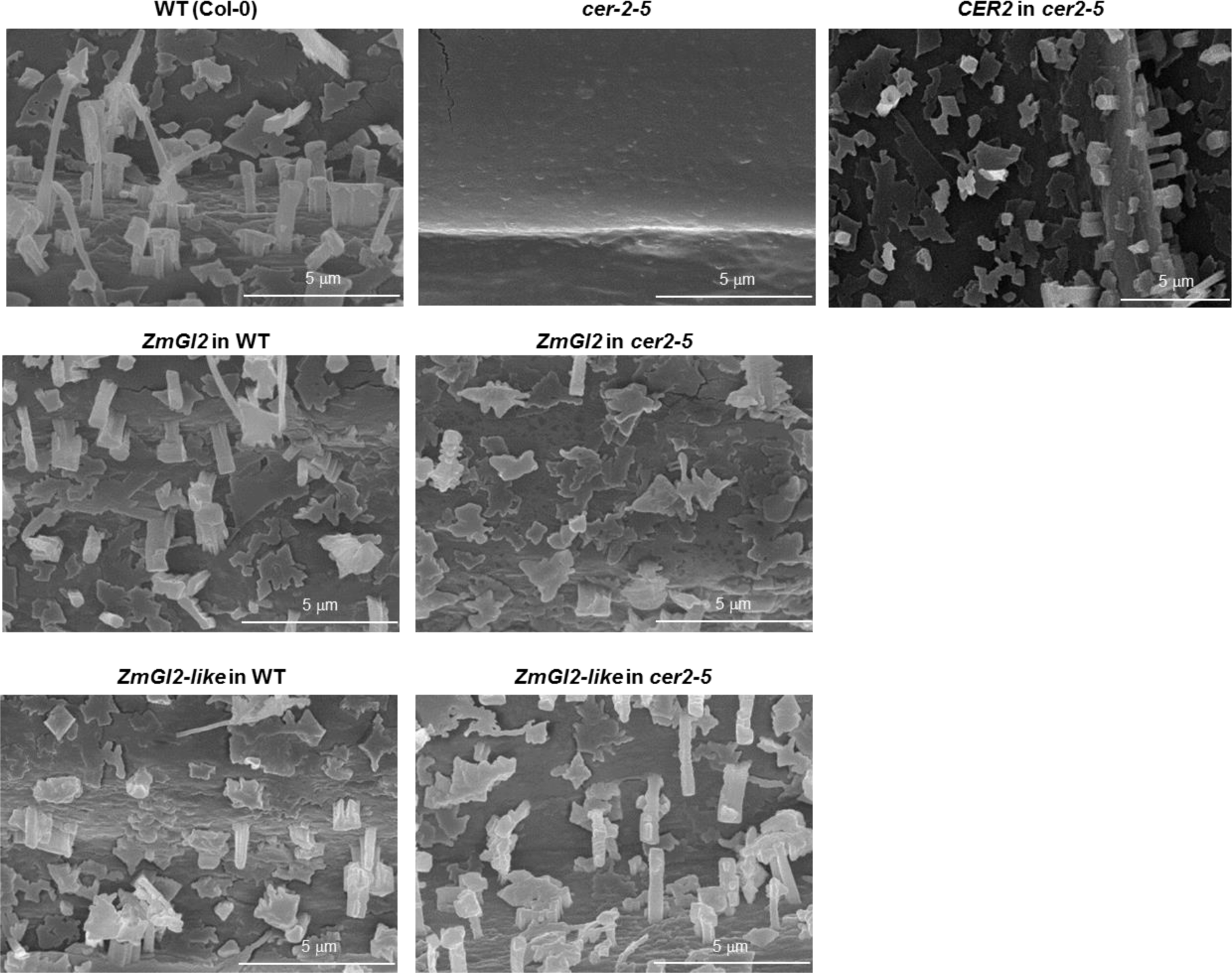
Arabidopsis stem epicuticular crystalloids. Scanning electron micrographs (10000X magnification) of stem surfaces of non-transgenic Arabidopsis wild-type or *cer*2-5 mutant plants, compared to stem surfaces of transgenic plants expressing *ZmGl2, ZmGl2-like or CER2* transgenes in either wild-type or *cer*2-5 mutant plants.

### Transgenic expression of ZmGL2 and ZmGL2-LIKE rescues the cuticular lipid-deficiency chemotype of the *cer2* mutant

Figure 4, Supplemental Figures 2 and 3, and Supplemental Table 1 present the data concerning the accumulation of extracellular cuticular lipids extracted from stems of the different genotypes developed in this study. The extractable cuticular lipid load on stems of the *cer2-5* mutant is about half of the wild-type stems (Supplemental Figure 2). As with the transgenic expression of *CER2,* the transgenic expression of either the maize *Gl2* or the *Gl2-like* gene in this mutant increases the cuticular lipid load by about two-fold, to near wild-type levels. The transgenic expression of *ZmGl2-like* in the wild-type background also increased the total cuticular lipid load by ∼15% compared to wild-type (statistically significant by t-test, p-value <0.05), whereas the expression of the *ZmGl2* transgene had no such effect on the total cuticular lipid load of the wild-type stems.

**Figure 4.**
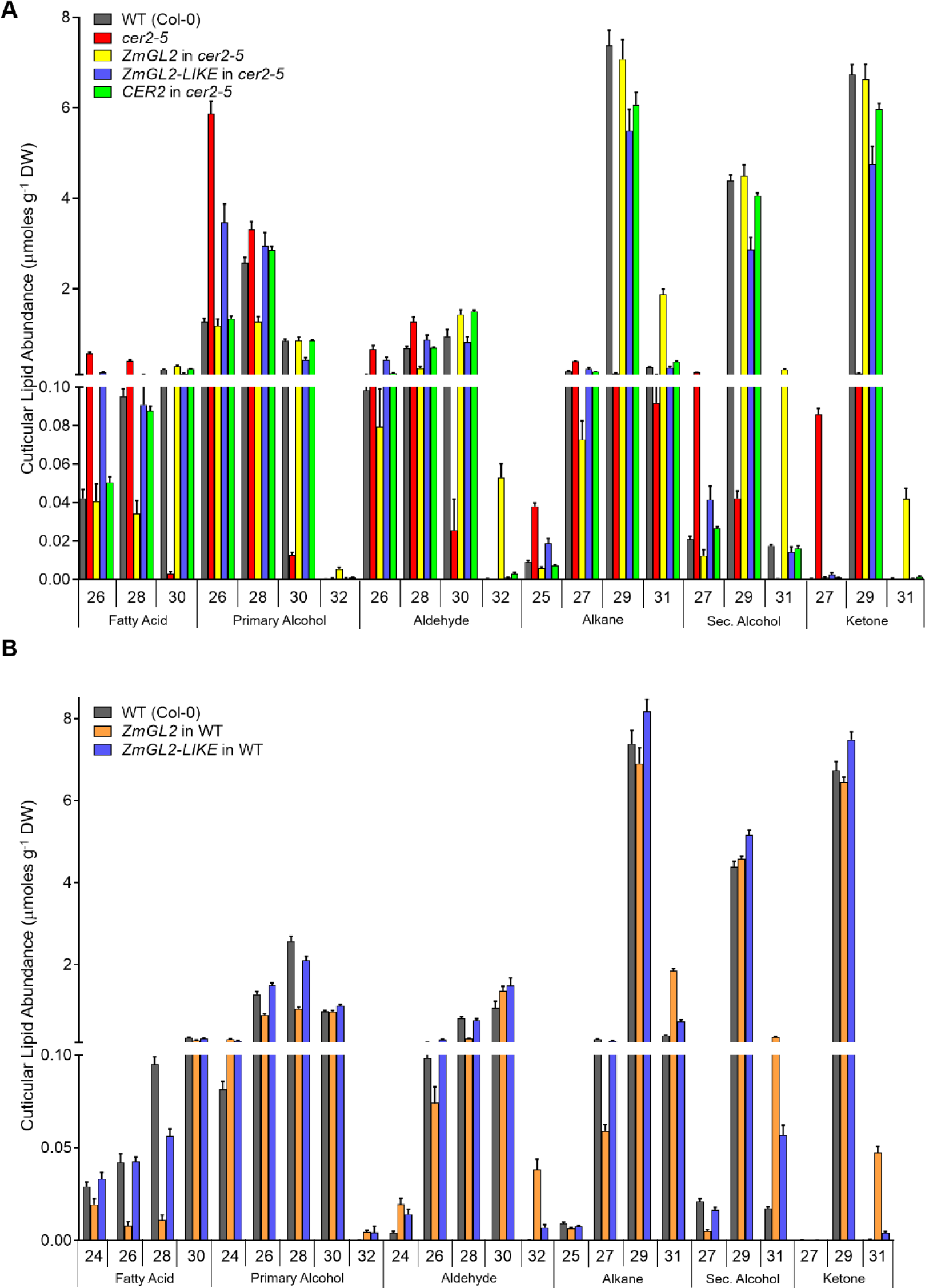
Effect of transgenic expression of *ZmGl2* and *ZmGl2-like* on the extracellular cuticular lipid profiles of Arabidopsis stems. A) Transgenic expression of *ZmGl2, ZmGl2-like or CER2* in the *cer2-5* mutant background, as compared to the profiles in the non-transgenic wild-type (Col-0) and *cer2-5* controls. B) Transgenic expression of *ZmGl2* and *ZmGl2-like* in the wild-type background. All numeric and statistical data, including the C20-C24 carbon chain-length minor components can be found in Supplemental Table 1. The data represent mean + standard error of 10-15 replicates

Figure 4 shows the compositional changes in the cuticular lipid profiles of these stems; these data are focused on alkyl chain-lengths of 26-carbons and longer, whereas the effect on the accumulation of the shorter length alkyl chains was negligible (these latter data are included in Supplemental Figure 2 and 3). The *cer2-5* mutation primarily affects the accumulation of the major components of the cuticular lipids, which are nearly eliminated in the mutant; these constituents are C30 fatty acid alkyl derivatives, specifically C29 alkane, C29 secondary alcohol, and C29 ketone. These decreases in accumulation are associated with a partial compensatory increase in shorter chain fatty acid derivatives, predominantly the C26 primary alcohol. The transgenic expression of either *Gl2*, *Gl2-like* or the *CER2* transgenes in the *cer2-5* mutant background results in a cuticular lipid compositional profile that is near identical to the wild-type profile in terms of the major cuticular lipid components. Thus, as compared to the *cer2-5* mutant, these transgenic lines show increased accumulation of the C29 alkane, C29 secondary alcohol, C29 ketone. However, the effect on the less abundant cuticular lipids is different among the three transgenes. Thus, like the *CER2* transgene, the *Gl2* transgene decreased the accumulation of the C26 components (i.e., primary alcohols) to wild-type levels, but the *Gl2-like* transgene was not capable of this effect, and the C26 primary alcohol remained at elevated levels as in the *cer2* mutant (Figure 4A).

In addition to the transgenic compensatory effects on the cuticular lipid profiles, the expression of the two maize transgenes in the *cer2-5* mutant induced novel changes to the profiles that are not normally present in wild-type controls. In particular, *ZmGl2* expression in *cer2-5* mutant induced the formation of longer alkyl chains; namely derivatives of the C32 fatty acid, which includes C31 alkane, C31 secondary alcohol, and C31 ketone. These novel metabolites are also observed when the *ZmGl2* transgene is expressed in the wild-type background (Figure 4). These latter novel components are not manifest by the transgenic expression of either the *CER2* or the *ZmGl2-like* genes in either the wild-type or the *cer2-5* mutant (Figure 4).

Collectively therefore, these biochemical changes establish that both the ZmGL2 and ZmGL2-LIKE proteins are functional homologs of CER2 and fully complement the biochemical deficiency associated with the *cer2* mutation. However, the two maize genes are not equivalent in how they affect cuticular lipid profiles; namely the *ZmGl2* transgene induces additional capabilities by enabling the production of even longer chain constituents than normal (up to 32-carbon fatty acids, and their alkyl derivatives), and the *ZmGl2-like* is incapable of reversing the effect on the accumulation of 26-carbon atom constituents.

### Mutations in the putative BAHD catalytic domain demonstrate differences in the *in vivo* **functionality between ZmGL2 and ZmGL2-LIKE**

The maize GL2 (Tacke et al., 1995), and the homologous Arabidopsis CER2 (Negruk et al., 1996; Xia et al., 1996) proteins initially defined the BAHD family of enzymes that catalyze acyltransferase reactions (D’Auria, 2006). A common structural feature is important to the catalytic function of BAHD enzymes, which has been identified through structural analysis and site-directed mutagenesis studies (Ma et al., 2005; D’Auria, 2006; Unno et al., 2007; Garvey et al., 2008). This is the HXXXDX-motif, which contains the catalytic His-residue that is responsible for deprotonating the alcohol or amine acyl acceptor-substrate, and is thus crucial to the acyltransferase catalytic mechanism. As with other biochemically characterized BAHD enzymes this motif is similarly positioned in the primary sequences of ZmGL2 and ZmGL2-LIKE proteins (Figure 1).

Interaction between potential substrates and this BAHD catalytic domain was computationally explored with structural models of ZmGL2 and ZmGL2-LIKE proteins, generated by *Phyre2* (Kelley et al., 2015). Using these structural models, the *3DLigandSite* algorithm (Kelley and Sternberg, 2009) identified the *Sorghum* hydroxycinnamoyl transferase (HCT) as the best BAHD structural template for ZmGL2 and ZmGL2-LIKE proteins. Using the experimentally determined structure of the substrate-enzyme complex of HCT, we identified that in addition to the potential catalytic His residue, the last “X” residue of the HXXXDX-motif of ZmGL2 and ZmGL2-LIKE (i.e., Ile-174 and Ile-190, respectively) may be sufficiently close to a potential substrate to directly interact. Furthermore, the Ile residue at this position is rare among BAHD homologs (occurs <1% of 1085 sequences that we analyzed), and residues with considerably smaller side chains are the most prevalent residues at this position (∼70% are Gly and ∼20% are Ala residues). As a contrast, the middle 3 “X” residues are more conserved as hydrophobic amino acids.

Therefore, site-directed mutagenesis was used to experimentally explore the functional role of the H, D and final “X” residues in the HXXXDX-domain of the ZmGL2 and ZmGL2-LIKE proteins. Specifically, with each protein we generated three point-mutants by substituting Ala for each of the three critical residues in this domain (i.e., GL2(H169A), GL2(D173A), GL2(I174A), and GL2-LIKE(H185A), GL2-LIKE(D189A) and GL2-LIKE(I190A)). Each of these mutants were transgenically expressed in the *cer2-5* mutant and wild-type Arabidopsis plants and the stem cuticular lipid profiles were analyzed to assess the ability of the mutant proteins to complement the *cer2-5* mutation (Figure 5, and Supplemental Table 2).

**Figure 5.**
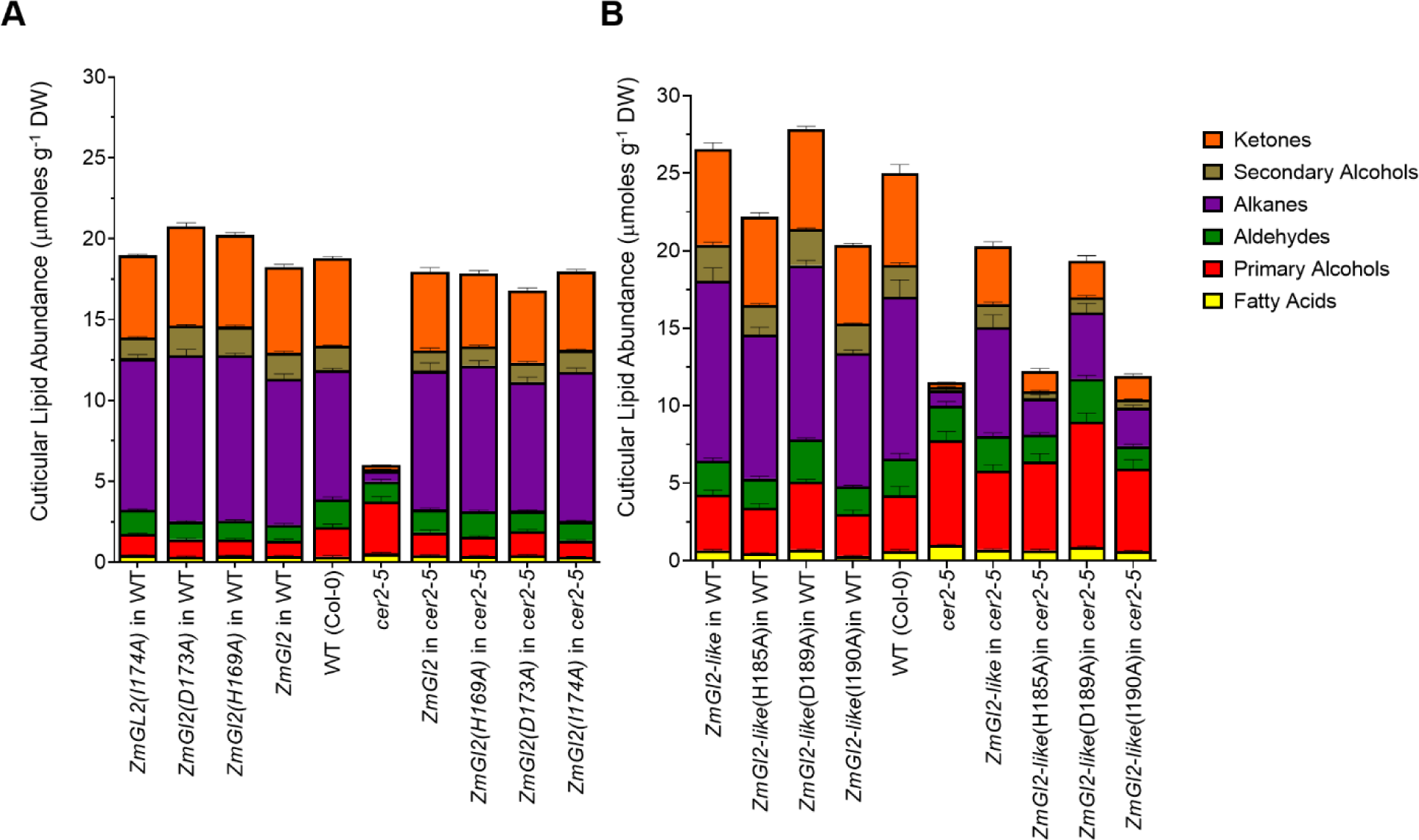
The role of the HXXXDX, BAHD-defining catalytic motif in supporting *in planta* functionality of ZmGl2 and ZmGl2-LIKE proteins. Transgenes were either the wild-type *ZmGl2* (A) or *ZmGl2-like* (B), or the indicated point mutants in the HXXXDX catalytic-motif. Each gene was expressed in Arabidopsis wild-type or *cer2-5* mutant genetic backgrounds. All numeric data and statistical analysis can be found in Supplemental Table 2. The data represent mean ± standard error of 10-40 replicates.

All three *ZmGl2* point mutants did not affect the ability of these transgenes to complement the *cer2-5* chemotype; namely either the expression of the wild-type *ZmGl2* transgene or any of the three *ZmGl2* point mutants, which should have destroyed the BAHD catalytic capability, are still capable of reversing the reduction in epicuticular lipid accumulation that is characteristic of *cer2-5* (Figure 5A). In contrast, the H185A, and I190A point mutants of *ZmGl2-like* transgene cannot fully complement the *cer2-5* chemotype, and reverse the cuticular lipid load on these stems (Figure 5B). These results demonstrate that the HXXXDX BAHD catalytic motif is not required for the ability of the ZmGL2 protein to functionally replace the CER2 function, however this motif is required for the ability of the ZmGL2-LIKE protein to fully replace the CER2 function.

### The ZmGL2 and ZmGL2-LIKE transgenes alter the VLCFA and hydroxy-VLCFA components of the cutin and cellular lipidome profiles

Because the genetic lesion that determines extracellular lipid traits occurs in the context of intracellular lipid metabolic processes, we profiled and compared the intracellular lipidomic pools in the stems of the different genotypes generated in this study. Furthermore, as a potential BAHD enzymes, ZmGL2 and/or ZmGL2-like maybe involved in the assembly of the ester bonds, which are prevalent in the assembly of the cutin polymer. For these reasons therefore, we also evaluated the effect of these genetic manipulations on the cutin monomers of the isolated cutin prepared from stems.

As with the cuticular lipid analyses described above, these comparisons are interpreted in the context of the effect of removing the *gl2* functionality in maize. Thus, lipidomics analyses indicate that mutation at the *gl2* locus in maize affects the VLCFA pool associated with the intracellular lipidome. These alterations are primarily associated with elongation of C26 and C28 fatty acids, leading to higher accumulation of these precursors (metabolites #60, #164 and #165, Supplemental Figure 4, Supplemental Table 3) and decreasing the level of C32 fatty acid (i.e., metabolite #167; Supplemental Figure 4, Supplemental Table 3).

Figures 6 illustrates the effect of the transgenic expression of *ZmGl2* (Fig. 6A) and *ZmGl2-like* (Fig 6B) on the accumulation of the three lipid classes, the cellular lipidome, cutin monomers and extractable cuticular lipids. These data indicate that quantitatively, the major effect of the *cer2* mutation is in halving the total accumulation of the extractable cuticular lipids, with minimal or no effect on the total accumulation of the cellular lipidome or cutin monomers. Moreover, the transgenic expression of *ZmGl2* or *ZmGl2-like* reversed the effect of the *cer2* mutation, and returned cuticular lipid content to wild-type levels, and had no quantitative effect on the total accumulation of the other lipid components that were evaluated (Figure 6A and B).

**Figure 6.**
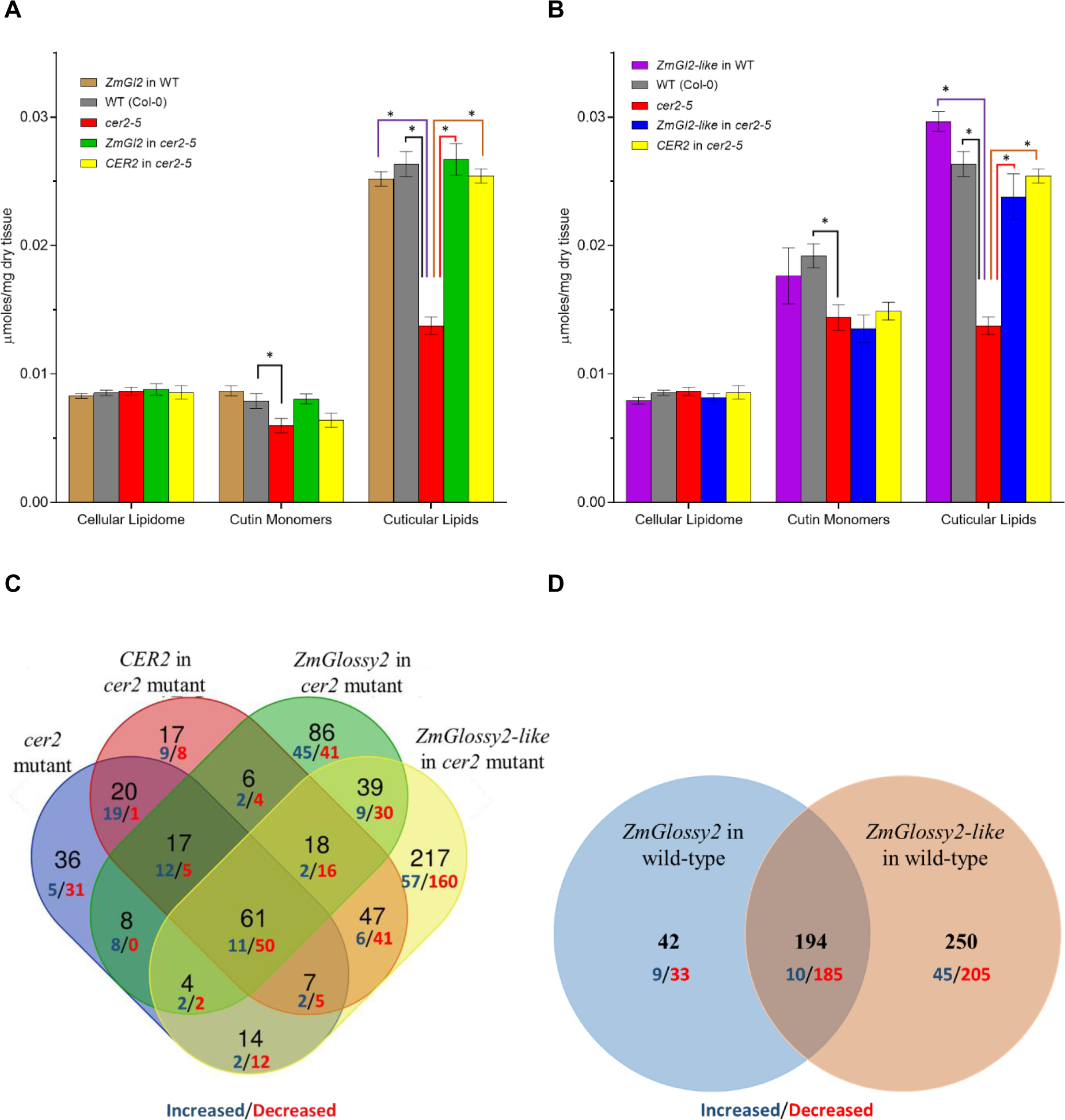
Effect of ZmGl2 and ZmGl2-like transgenes on accumulation of Arabidopsis stem lipids. Comparison of the cellular lipidome, cutin monomer and cuticular lipid content of non-transgenic wild-type and *cer2-5* mutant stems and transgenic plants expressing *ZmGl2* or *CER2* (A) and *ZmGl2-like* or *CER2* (B). The data represent mean ± standard error of 4-15 replicates. Asterisks indicate genetic manipulations that result in statistically significant changes as evaluated by Student’s T-test (p-value & 0.05). Venn-diagram representation of changes in the accumulation of analytes identified in the cellular lipidome of stems in response to the transgenic expression of *ZmGlossy2*, *ZmGlossy2-like* and *CER2* in the *cer2-5* mutant background (C) or in a wild-type background (D). Each subset identifies the number of lipid analytes whose abundance is either increased (blue digits) or decreased (red digits) in the indicated genetic background. The digital data can be found in Supplemental Table 3. The altered accumulation levels are statistically significant based on evaluation by Student’s T-test (p-value & 0.05).

However, these transgenic manipulations had compositional effects on these lipids, and we focus the following text on the effect on VLCFA and derivative components. Although the cutin polymer is thought not to have VLCFA components, cutin preparations often contain VLCFA derivatives (i.e., 2-hydroxy-VLCFAs), which are probably associated with sphingolipids that co-purify with cutin (Molina et al., 2006). Thus, regardless of their complex-lipid origins, because these VLCFAs are FAE-generated products, we evaluated the effect of *ZmGl2* and *ZmGl2-like* transgenic expression on their abundance.

Specifically, in the *cer2-5* mutant there are reductions in accumulation of the C22- and C24-fatty acids and their 2-hydroxy-derivatives, and an increase in the C26 fatty acid and the corresponding 2-hydroxy-derivative that was associated with the cutin preparations (i.e., metabolite #163, #165, #167, #176, #178, and #181, Supplemental Figure 5A, Supplemental Table 3). Transgenic expression of *Gl2* in the *cer2-5* mutant reversed these alterations in VLCFA components, and in some cases further enhanced their accumulation to double or quadruple the levels that occur in the wild-type controls (i.e., metabolites #163, #165, #176, #178, and #179, Supplemental Figure 5C). In contrast, the transgenic expression of *Gl2-like* caused only minor alterations in the total levels of cutin monomers (Fig. 6B) and of the VLCFA components associated with the cutin preparations, and this was irrespective of whether *Gl2-like* is expressed in a wild-type or *cer2-5* mutant background (Supplemental Figure 5E and 5F, Supplemental Table 3).

Insights into the effect of transgenic expression of *ZmGl2* and *ZmGl2-like* on the cellular lipidome was provided by the analysis of lipid extracts via a standardized LC-qTOF analytical platform (Okazako and Saito, 2018). These analyses determined the relative abundance of 1372 lipid analytes, 153 of which were chemically identified. The latter include phosphoglycerolipids (29), glycolipids (35), isoprenoids (24), acylated glycolipids (8), sterol lipids (12), storage lipids (33) and free fatty acids (12) (Supplemental Table 3). Among these chemically defined lipids, the most striking alterations are the changes in the accumulation of the free fatty acids (i.e., metabolites #148 to #153, Supplemental Figure 5A-F, Supplemental Table 3). The *cer2-5* mutation reduced the accumulation of C30 fatty acid (metabolite #150), resulting in the higher accumulation of the precursor fatty acids of 26 and 28 carbon chain lengths (i.e., metabolites #148 and #149, Supplemental Figure 5A, Supplemental Table 3). The transgenic expression of maize *Gl2* and *Gl2-like* in this mutant reversed these *cer2-5*-induced effects (Supplemental Figure 5C and 5E, Supplemental Table 3). Moreover, *ZmGl2* has additional capabilities, inducing the increased accumulation of C32 fatty acid (i.e., metabolite #151, Supplemental Figure 5C and 5D, Supplemental Table 3). This latter effect correlates with the increased levels of C31 alkyl derivatives that were detected in the epicuticular lipid profiles of wild-type and *cer2-5* mutant stems in the same transgenic lines (Figure 4). These findings suggest that both *ZmGl2* and *ZmGl2-like* have the ability to affect the terminal elongation processes of fatty acids to C30 and C32 chain-lengths, which are negatively impacted by the *cer2* mutation.

Another alteration in the lipidome that is consistent with the shared functionality between *CER2, ZmGl2* and *ZmGl2-like* in fatty acid elongation is the alteration in the accumulation pattern of the glycosylceramides that utilize a 2-hydroxy C26 fatty acid building block (i.e., metabolites #58 and #59, Supplemental Figure 5A, Supplemental Table 3). The accumulation of these metabolites is doubled in the *cer2-5* mutant, and this effect is reversed by the transgenic expression of either *ZmGl2* or *ZmGl2-like* (Supplemental Figure 5C and 5E, Supplemental Table 3). An additional similar genetic modulation, but not associated with fatty acid elongation, is the accumulation of the minor sterol esters; i.e., β-sitosterol esterified with linoleic or linolenic acid (i.e., metabolites #106 and #107, Supplemental Figure 5A, 5C and 5E, Supplemental Table 3).

Compared to the wild-type, the accumulation of these two sterol esters is increased by between 10 and 25 fold in the *cer2-5* mutant, and their levels are decreased to normal when *ZmGl2* is transgenically expressed in the mutant; in contrast, the transgenic expression of *ZmGl2-like* does not have this latter effect (Supplemental Figure 5C and 5E, Supplemental Table 3).

These lipidomics analyses also quantified changes in the accumulation of 1,219 analytes that are chemically undefined (Supplemental Table 3). On a mole basis these chemically undefined lipids account for ∼15% of all the lipids that were collectively profiled in the wild-type and in the *cer2-5* mutant tissue. These data would be more informative of *ZmGl2* or *ZmGl2-like* functions once their chemical identities are determined, however, we evaluated their relative accumulation patterns as molecular markers to gain insights into the relative functionalities of the two maize genes. In the tabular data presented in Supplemental Table 3, these analytes are identified by the *m/z* values of the individual ions detected by LC-qTOF in either negative or positive ion modes.

The accumulation of approximately half of these chemically undefined lipid analytes are unaffected by the *cer2*-associated genetic manipulations. The Venn-diagram shown in Figure 6C illustrates the relative abundance of the other half of these undefined analytes, which can be classified into 15 categories based on the change in their relative abundance in response to the genetic manipulations generated in this study. Specifically, 35% of these analytes showed decreased accumulation in response to these genetic manipulations, whereas only 15% showed increased accumulation. As molecular markers, the changes in the accumulation of these unknown metabolites associated with transgenic expression of *ZmGl2* or *ZmGl2-like* do not parallel the genetic complementation displayed by *CER2* trans-expression indicating that the three genes are not equivalent in complementing the *cer2* mutant trait (Figure 6C). For example, there are 156 analytes (13% of detected analytes), whose accumulation is similarly altered by the expression of either *CER2*, *ZmGl2* or *ZmGl2,* but in contrast there are 17, 86 and 217 analytes (collectively 26% of the detected analytes) that are uniquely altered by the expression of each individual transgene. Moreover, the expression of the two maize homologs in a wild-type background induce changes in approximately 300 analytes of which only 2/3^rd^ are common to both transgenic lines (Fig 6D). Therefore, although the two maize homologs can functionally complement the cuticular lipid phenotype of the *cer2* mutation, the two maize genes also express unique attributes that are not equivalent to each other or to the *CER2* gene.

## DISCUSSION

The extracellular cuticular lipids that coat the outer surfaces of the aerial organs of terrestrial plants (Fernandes et al., 1964; Kolattukudy 1980) are specifically produced by the epidermal cell layer of these organs. Because epidermal cells account for only about 10% of the cellular population of these aerial organs (Jellings and Leech, 1982), elucidating the molecular and biochemical mechanisms that regulate their biogenesis is confounded by the other 90% of the cell population that is not involved in these processes. This technical barrier has been partially overcome by utilizing forward genetic approaches to identify and characterize genes involved in extracellular cuticular lipid accumulation (Kunst and Samuels, 2009; Shaheenuzzamn et al., 2019). This strategy has been successful in the isolation of causative genes that generate the cuticular lipid phenotypes (*glossy* in maize and *eceriferum* in Arabidopsis). Yet in many cases, the exact mechanisms by which these gene products affect cuticular lipid deposition is still unclear.

Exemplary of such molecular characterization of cuticular lipid genes is the *gl2* gene of maize, and its Arabidopsis homolog, *cer2* (Tacke et al., 1995; Xia et al., 1996). In this study we specifically implemented a transgenic strategy to characterize the functional inter-relationship between the maize *Gl2* gene, and the homologous *Gl2-like* gene, and the Arabidopsis homolog, *CER2*. We identified the maize *Gl2-like* gene by the shared 63% sequence similarity with *Gl2*. Although *ZmGl2* is known to be involved in cuticular lipid biosynthesis (Hayes and Brewbaker, 1928; Bianchi, 1975; Tacke et al., 1995*), ZmGl2-like* may have overlapping but distinct roles in this pathway. The transgenic experiments conducted in this study utilized Arabidopsis as the vehicle for these evaluations, testing for the ability of each maize gene to compensate for the missing function associated with the homologous Arabidopsis *CER2* gene; in parallel we also evaluated the effect of each transgenic event in the wild-type background.

### The ability of *Gl2* and *Gl2-like* to contribute to extracellular cuticular lipid deposition

The functionality of *ZmGl2* in cuticular lipid biosynthesis was initially identified by the genetic observations that mutations at this locus eliminate the deposition of these constituents on the surfaces of juvenile seedling leaves (Hayes and Brewbaker, 1928; Bianchi, 1975). The identification of *ZmGl2-like* is solely based on sequence homology with the ZmGL2 and CER2 proteins, without any functional data to support its role in cuticular lipid biosynthesis. Mutations at the *cer2* locus block the normal accumulation of cuticular lipids, by apparently blocking the conversion of C26 and/or C28 fatty acid to C30 fatty acid (Mcnevin et al., 1991; Jenks et al., 1995). Our transgenic studies established the functionality of *ZmGl2-like* in cuticular lipid biosynthesis. Specifically, as with the transgenic expression of *CER2* in a *cer2* mutant, the transgenic expression of either *ZmGl2* or *ZmGl2-like* in the same mutant, restores the normal accumulation of cuticular lipids, and restores the ability to convert the C26/C28 fatty acids to C30 fatty acid. Therefore, both *ZmGl2* and *ZmGl2-like* are functional homologs of *CER2*. However, there are differences induced by the transgenic expression of *ZmGl2* as compared to *ZmGl2-like*, which suggest that the two gene products have acquired distinct functionalities since the probable gene duplication event that gave rise to the two homologs. These differences between *ZmGl2* and *ZmGl2-like* are revealed as differences in the 1) ability to complement the *eceriferum* phenotype on Arabidopsis stems; 2) epicuticular crystalloid morphologies; 3) total cuticular lipid loads and compositions; and 4) intracellular lipidomes in the *cer2-5* lines complemented by each maize gene. In these comparisons, the *cer2* plants that are genetically complemented with the *ZmGl2-like* transgene present traits that are more like the wild-type Arabidopsis plants than the *ZmGl2*-complemented plants. Moreover, although the transgenic expression of either the *CER2* gene or *ZmGl2-like* in the wild-type background did not induce any new cuticular lipid components, the parallel experiments conducted with the transgenic expression of *ZmGl2* result in plants that express new cuticular lipid components that are not associated with the wild-type Arabidopsis host (i.e., the ability to support the generation of C32 fatty acids and alkyl derivatives). Collectively, these observations are interpreted to indicate that although both maize *Gl2* and *Gl2-like* genes express a functionality that can replace the *CER2* function of Arabidopsis, the *ZmGl2-like* gene more completely and accurately recapitulates the *CER2* functionality, and *ZmGl2* encodes additional functionalities that are beyond those of *ZmGl2-like* and *CER2* genes. This neofunctionalization that probably arose following the gene duplication event that generated the two maize paralogs enables *ZmGl2* to induce the production of C32 alkyl-chains in Arabidopsis is consistent with the fact that the maize cuticular lipids are predominantly alkyl chains derived from C32 fatty acids.

### Relationship between BAHD catalytic activity and the ability of *Gl2* and *Gl2-like* to support ***in planta* epicuticular lipid deposition**

Although not initially recognized (Tacke et al., 1995; Xia et al., 1996), CER2 and ZmGL2 proteins proved to be archetypal of the BAHD-family of acyltransferases (St-Pierre and Luca, 2000; D’Auria, 2006). This catalytic ability of the BAHD enzymes was demonstrated by the biochemical characterization of a number of enzymes that catalyze acyltransferase reactions in the biosynthesis of diverse specialized metabolites. The BAHD enzymes are recognizable by the conservation of two primary sequence motifs: a) the HXXXDX catalytic domain; and b) the DFGWG domain that appears to stabilize the substrate-enzyme complex (St-Pierre and Luca, 2000; D’Auria, 2006). Although both domains have been reported to be essential for enzyme functionality, the latter domain is not fully conserved among all characterized BAHD enzymes (Suzuki et al., 2003; Bayer et al., 2004; Ma et al., 2005; Unno et al., 2007).

The neofunctionalization that was enabled by the gene duplication that gave rise to *ZmGl2* and *ZmGl2-like* appears to be associated with the potential catalytic capabilities of this BAHD-defining catalytic domain. The role of the HXXXDX domain to support cuticular lipid deposition by ZmGL2 and ZmGL2-LIKE were tested *in planta* by directed mutagenesis. Point mutants that are predicted to disrupt BAHD catalytic activity (i.e., the His, Asp and terminal “X” residue of the HXXXDX domain) did not affect the ability of ZmGL2 to support cuticular lipid deposition, whereas these mutations partially disrupted the ability of ZmGL2-LIKE to support this metabolic outcome. These findings indicate that although the ZmGL2 protein does not require BAHD catalytic activity to support cuticular lipid deposition, this domain is required for GL2-LIKE to fully achieve the same function.

The ZmGL2 results agree with studies previously conducted with CER2, which demonstrated that the BAHD-defining HXXXDX domain is not needed to complement the *cer2*-associated cuticular lipid deficiency (Haslam et al., 2012). Moreover, this study also demonstrated that the yeast co-expression of CER2 with a 3-ketoacyl-CoA synthase (KCS) component of the plant FAE system results in the ability of the yeast strain to produce longer-chain fatty acids than normal, up to 28 and 30 carbon chain lengths; yeast strains normally only produce C26 VLCFA. Moreover, this latter capability does not require an intact HXXXDX catalytic domain. These results have been interpreted to indicate that CER2 non-catalytically interacts with the FAE system for VLCFA biosynthesis, and alters the product profile of the FAE system. Similar conclusions have been reached with the characterization of Arabidopsis *CER2* homologs, such as *CER2-LIKE1* and *CER2-LIKE2*, where the former lacks the His residue in the predicted HXXXDX motif (Haslam et al., 2012, 2015).

In contrast to *CER2* and *ZmGl2, ZmGl2-like* needs the intact, and presumably functional, BAHD-defining HXXXDX catalytic domain to fully complement the *in planta cer2* function. Moreover, a functional HXXXDX, BAHD-catalytic domain appears to be required by the rice *CER2* homolog, *WSL4*, which facilitates the interaction with KCS (Wang et al., 2017), to affect fatty acid elongation beyond C24 chain-lengths. These characterizations raise a number of questions concerning the ability of these proteins to support cuticular lipid deposition and the function of other Clade II BAHD proteins: 1) is ZmGL2-LIKE (and possibly the rice homolog, OsCER2) a bifunctional protein, one mediated by the HXXXDX catalytic function, and the second that is independent of this catalytic function; 2) do other members of the Clade-II BAHD family possess an acyltransferase catalytic activity; and 3) what are the two substrates (the acyl-donor and acyl-acceptor substrates) of the ZmGL2-LIKE acyltransferase enzyme. Answering these questions will probably require the development of a direct biochemical assay that goes beyond the studies conducted to date; these latter studies primarily rely on correlations between genetic modifications and metabolic outcomes.

### The role of *Gl2* and *Gl2-like* in supporting the accumulation of VLCFA components of intracellular lipid pools

Mutations in the *cer2* locus appear to block the fatty acid elongation process from C26 or C28 to C30 fatty acids (McNevin et al., 1993; Jenks et al., 1995). Based on our current understanding of the mechanism of fatty acid elongation, it is mechanistically unclear how specific iterations of the elongation cycle (i.e., from C26 or C28 to C30) can be distinguished from any of the other six iterations that elongate a C18 fatty acid to a C30 fatty acid. Recent studies with the Arabidopsis CER2 and rice homologs, have proposed the possibility of a physical interaction between the condensing enzyme of the FAE complex (specifically KCS6) and CER2 (Haslam et al., 2015; Wang et al., 2017) and this interaction may affect a single iteration of the elongation cycle.

Based on the cuticular lipid profiles in transgenic Arabidopsis lines we surmise that as with *CER2, ZmGl2-like* can support the ability of elongating fatty acids from C28 to C30. Although *ZmGl2* also shares this capability, it can additionally contribute to the elongation of C30 to C32 fatty acids. Thus, we hypothesize that *ZmGl2-like* and *ZmGl2* probably have overlapping functions in maize, where the former affects fatty acid elongation from C26/C28 to C30, and the latter the elongation from C26/C28 to C32.

The primary genetic lesion that determines these extracellular cuticular lipid traits occur in the intracellular lipid metabolic processes that underlie the origins of the cuticular lipids. These processes are primarily associated with the intracellular membranes (ER and possibly plasma membrane), which house the biochemical reactions associated with VLCFA biosynthesis and down-stream reactions that generate the other alkyl derivatives of the cuticle (Bernard and Joubès, 2013). By profiling and comparing the intracellular lipidomic pool and the cutin matrix of the stems of the different genotypes generated in this study, we evaluated the extent to which VLCFA metabolism is juxtaposed with other lipid metabolic pathways (e.g., phospholipids, sphingolipids, and storage lipids). These comparisons indicate that genetic manipulations associated with mutating the *cer2* function and replacing it with either *ZmGl2* or *ZmGl2-like*, not only affects cuticular lipid profiles, but they also have pronounced effects on those lipids that utilize VLCFA building blocks (Li-Beisson et al., 2010; Bernard and Joubès, 2013). In addition, *CER2, ZmGl2,* and *ZmGl2-like* altered the accumulation of several unidentified intracellular lipid molecules. In combination therefore, these data imply that these genes may have regulatory roles in controlling the FAE complex.

Based on the lipidomics analyses and the HXXXDX-mutagenesis studies, it is possible to hypothesize that the acyltransferase reaction of ZmGL2-LIKE may be part of the regulatory mechanism that modulates the FAE complex, particularly the terminal elongation cycle. However, it’s unclear how an acyltransferase reaction mechanism can affect one of several FAE reaction cycles that utilize iterations of Claisen condensation-reduction-dehydration-reduction reaction mechanisms.

## MATERIAL AND METHODS

### Plant Material and Growth Conditions

T-DNA mutant line SALK_084443C (*cer2-5*; At4g24510) in the Col-0 background was obtained from the Arabidopsis Biological Resource Center (www.arabidopsis.org). This T-DNA insertion disrupts the second exon of *CER2.* This *cer2-5* mutant stock and the wild-type Col-0 stock were used in all the experiments described herein. Seeds were sown in LC1 Sunshine Mix (Sun Gro Horticulture, Bellevue, WA) and treated at 4°C for 5d to break seed dormancy. Seedlings were transferred to individual pots and grown to maturity under constant growth conditions in a regulated growth room at 22°C under continuous illumination (2568 Lux or photosynthetic photon flux density of 100 μmol of photons m^-2^ sec^-1^). Plants were watered once a week and the irradiance, temperature, and relative humidity were monitored using an Onset Computer Corporation (Bourne, MA) Hobo monitor U12-012 (www.onsetcomp.com). Biochemical and microscopic analyses were conducted on stem tissues of Arabidopsis plants of different genotypes, harvested when the primary flower bolt was 35 to 40-cm in height. Flowers, cauline leaves and siliques were removed and stem tissue was used for analysis.

Maize *glossy2* mutant seeds (*glossy2-Salamini*; Maize Genetics COOP Stock Center catalog #208H; maizecoop.cropsci.uiuc.edu) were out-crossed to inbred B73 and the resulting heterozygous F1 seeds were selfed and back-crossed to the inbred B73. The BC1 seeds were planted in soil, and grown in a climate-controlled greenhouse under a diurnal cycle of 16-h light and 8-h dark at 27°C and 24°C respectively, maintaining 30% relative humidity. The *gl2* mutant seedlings were identified from the segregating progeny by their water beading phenotype, and non-beading sibling seedlings were used as wild-type controls. Five individual maize seedling plants, at the 3-5 leaf stage were pooled to generate a single replicate, and a total of 4 to 5 replicates were used for cuticular lipid and lipidomic analyses.

### Molecular Cloning

The full-length *Gl2* (GRMZM2G098239; Zm00001d002353) and *Gl2-like* (GRMZM2G315767; Zm00001d024317) ORFs were codon-optimized for expression in Arabidopsis with GeneOptimizer (GeneArt, LifeTechnologies) and OptimumGene^TM^ (GenScript, Piscataway, NJ; www.genscript.com), respectively, and these sequences were chemically synthesized (GenScript, Piscataway, NJ; GeneArt, LifeTechnologies). The *ZmGl2* sequence was cloned into pMA-RQ and pDONR221 entry vectors by Life Technologies Corporation (Carlsbad, CA, USA). *ZmGl2-like* sequence was initially obtained from GenScript as a pUC57-clone and was sub-cloned into pENTR^TM^/D-TOPO^®^ entry vector (Invitrogen). Further sub-cloning of *ZmGl2* and *ZmGl2-like* sequences for plant expression experiments were performed using LR Clonase II Enzyme Mix (Invitrogen), using pEarleyGate100 vector (Earley et al., 2006), which controls expression of the transgene with the CaMV 35S promoter. The resultant recombinant vectors (p35S::*ZmGlossy2* and p35S::*ZmGlossy2-like*) were introduced into *Agrobacterium tumefaciens* (strain C58C1) by electroporation (Sambrook et al., 1989). For site-directed mutagenesis experiments, mutation of the His-, Asp-, and Ile-residues of the HXXXDX motif of ZmGL2 and ZmGL2-LIKE were generated using QuikChange XL Site-Directed Mutagenesis Kit (Agilent Technologies, Santa Clara, CA). The ZmGL2 protein was also expressed in *E. coli* BL21AI strain (Invitrogen, Carlsbad, CA, USA, www.thermofisher.com) using the pDEST17 expression vector (Invitrogen). The authenticity of all recombinant plasmids was confirmed by DNA sequencing.

### Plant Transformation and Selection

*ZmGl2* and *ZmGl2-like* transgenes were transformed into wild-type Arabidopsis, Col-0 ecotype, and the homozygous *cer2-5* mutant line via an Agrobacterium-mediated floral-dip protocol, adapted from Clough and Bent, 1998. Briefly, inflorescence bolts were submerged for 20s in an infiltration medium containing *A. tumefaciens* (strain C58C1) carrying either p35S:*ZmGlossy2* or p35S::*ZmGlossy2-like.* The infiltration medium consisted of 2% sucrose and 0.02% Vac-in-stuff Silwet L-77 (Lehle Seeds, Round Rock, TX). Plants were then returned to a 22°C growth chamber under continuous illumination until seed-set. Seeds from transformed plants were collected, and germinated in soil. Transformed seedlings were initially identified as being resistant to BASTA herbicide (i.e., glufosinate), applied at a dilution of 1:1000. The herbicide resistant plants were propagated and selfed to the T2 generation. Plants carrying the *Gl2* or *Gl2-like* transgene were molecularly confirmed with gene-specific primers by PCR analysis using DNA-templates isolated from individual herbicide resistant plants. RNA expression of *ZmGl2-like* was confirmed by RT-PCR and ubiquitin mRNA (At4g05320) was used as the internal control, GL2 protein expression was evaluated by Western blot assays. Three replicate lines from each of three independent transformation events were maintained and used for all further experiments.

### Western blot analysis

Protein-extracts were prepared by homogenizing plant leaf tissue with a buffer consisting of 62.5 mM Tris-HCl, pH 6.8, 30% glycerol, 10% SDS and 10% 2-mercaptoethanol. Samples were vortexed for 5-min, boiled for 10-min and centrifuged at 13000g for 2-min. The clarified supernatant protein extracts were subjected to SDS-PAGE and proteins were transferred to a nitrocellulose membrane according to manufacturer’s instructions (Bio-RAD, Hercules, CA). Protein blots were first probed with GL2-specific antibody, recovered from ascites fluid recovered from GL2 challenged mice (www.biotech.iastate.edu/Hybridoma) (1:1000 dilution), and then with horseradish peroxidase-linked anti-mouse IgG antibody (Bio-Rad) (1:3000 dilution). The antigen-antibody complexes were detected using the Pierce ECL chemi-luminescent detection system (Thermo Scientific, Rockford, IL) and visualized on the ChemiDoc XRS+ gel documentation system (Bio-Rad).

### Stereomicroscopy

The *eceriferum* phenotype was visualized from the central 1-cm segment of the stems using a Zeiss macro-Zoom microscope (Zeiss Axio Zoom V16) with ZEN2 software (Carl Zeiss Inc., Thornwood, NY).

### Scanning Electron Microscopy

The 1-cm-long stem segments were mounted on aluminum stubs with double-sided carbon tape, dried in a desiccator and sputter-coated (www.tedpella.com) with a Cressington HR208 sputter coater with platinum for 90s at 40 milliamps, depositing a 10 nm-thick coating. The segments were examined at 10 kV with a Hitachi SU4800 field emission SEM (www.hitachi-hightech.com), and images were digitally captured in TIFF format.

### Extraction and analysis of stem extracellular cuticular lipids

Extracellular cuticular lipids were extracted from isolated tissue by immersing the stems for 10-seconds in 10-mL chloroform (HPLC-grade Fisher Chemicals, Pittsburgh, PA), containing 5-µg hexacosane (Sigma-Aldrich, Milwaukee, WI) as an internal standard. The stems were then flash frozen using liquid nitrogen, lyophilized (Labconco FreeZone Benchtop Freeze Dry System), and the weight of the dry biomass was determined. A similar extraction procedure was used to extract cuticular lipids from maize seedling, collected at 3-5 leaf stage, using 60-μg hexacosane as the internal standard.

The chloroform was removed from the extracellular cuticular lipid extracts by evaporation under a stream of N_2_ gas, and the dried lipids were silylated using a protocol based on that of Wood et al., (2001) and Hannoufa et al., (1993). Specifically the dried extracts were dissolved in 0.2-mL acetonitrile and silylation was performed by the addition of 0.05-mL of *N*,*O*-Bis(trimethylsilyl)trifluoroacetamide (BSTFA) with 1% trimethylchlorosilane (TMCS) (Sigma-Aldrich), and incubated at 65°C for 30 min. The samples were cooled, dried under a stream of N_2_ gas and resuspended in chloroform. One-microliter of the derivatized sample was injected into GC-MS or GC-FID.

GC-MS analysis was conducted with an Agilent 7890A GC, equipped with a HP-5ms capillary column (30 m x 0.25 mm x 0.25 μm, Agilent), interfaced to an Agilent 5975C quadrapole mass spectrometer. Chromatography was conducted with helium gas, at a flow-rate of 1.0 mL/min, and an inlet temperature at 280°C. The column oven temperature was initially held at 120°C, then ramped at 10°C/min to 260°C and held at this temperature for 10 min, and then ramped at 5°C/min to 320°C and held there for 4 min. EI-MS ionization energy was set to 70 eV and the interface temperature was 280°C. Resulting chromatograms and mass-spectra were deconvoluted and queried against an in-house Mass-Spectral library and the NIST 14 Mass Spectral Library using the NIST AMDIS software (Stein, 1999).

GC-FID analysis was conducted with an Agilent 6890 GC, equipped with a DB-1 MS capillary column (15 m x 0.25 mm x 0.25 μm, Agilent 122-0112). Chromatography was conducted with helium gas, at a flow-rate of 1.2 mL/min, and an inlet temperature at 280°C. The column oven temperature was initially held at 80°C, then ramped at 15°C/min to 220°C, then ramped at 7.5°C/min to 310°C, and finally ramped at 20°C/min to 340°C and held for 5 min. The Agilent ChemStation software was used for peak alignments, and parallel GC-MS analysis was used to chemically identify eluting peaks.

The relative abundance of extracellular lipids/mg dry weight of plant material was calculated based on the ion-strength of the internal standard. Statistical significance was determined using Student’s t-test.

### Extraction and analysis of stem cutin monomers

Using procedures described by Li-Beisson et al., (2013), cutin was extracted from 50-100 mg of freshly harvested tissue, spiked with 10-μg heptadecanoic acid as an internal quantitation standard. The monomer components were converted to methyl esters by acid-catalysis and analyzed by GC-MS methods as described by Li-Beisson et al., (2013).

### Extraction and analysis of stem cellular lipidome

Intracellular lipids were extracted from 5-8 mg dry Arabidopsis stem tissues or dried maize seedling leaf tissues that had previously been extracted for epicuticular lipids. The samples were analyzed using a Waters Xevo G2 Q-TOF MS equipped with a Waters ACQUITY UPLC system as previously reported by Okazaki et al., 2018. Briefly, flash frozen plant tissue was lyophilized and pulverized to fine powder using a Mixer Mill 301 (Retsch GmbH, Germany). An extraction solvent containing a mixture of methyl tert-butyl ether and methanol (3:1, v/v), spiked with 1 μM 1,2-didecanoyl-sn-glycero-3-phosphocholine (Sigma-Aldrich) as internal standard, was added to the tissue, at a rate of 160 μl per mg dry tissue and the mixture was thoroughly vortexed. Water was added (50-μl per mg tissue) and the mixture was thoroughly mixed for 10-min at room temperature using a sample tube mixer. After a 10-minute incubation on ice the mixture was centrifuged at 3000 *x* g at 4°C for 10 min. 160 μl of the supernatant was collected and evaporated to dryness in a SpeedVac. The residue was dissolved in 200 μl of ethanol, vortexed for 10-min at room temperature, and centrifuged at 10,000 g at 4°C for 10-min. The supernatant (180-μl) was used immediately for lipid analysis. Lipid analysis was conducted using liquid chromatography quadrupole time-of-flight mass spectrometer (HPLC, Waters Acquity UPLC system; MS, Waters Xevo G2 Qtof), as reported by Okazaki et al., 2018.

The relative abundance of the cellular lipidome was calculated using abundances detected in positive ion mode for all lipid categories (including unknowns) with the exception of fatty acids, which were detected in negative ion mode. Statistical significance was determined using Student’s t-test.

## Supplemental Material

**Supplemental Table 1. Cuticular lipid abundance of Arabidopsis stems and maize leaves**

**Supplemental Table 2. Cuticular lipid abundance of Arabidopsis stems expressing mutated *Gl2* and *Gl2-like* transgenes**

**Supplemental Table 3. Lipidome and cutin data**

## ACKNOWLEDGEMENTS

The authors acknowledge Dr. Harry T. Horner, Tracey Stewart and Randell Den Adel for helpful discussion and for use of equipment at Roy J. Carver High Resolution Microscopy Facility (Iowa State University, Ames, IA); Drs. Zhihong Song, Ann Perera and Lucas Showman of the WM Keck Metabolomics Research Laboratory (Iowa State University, Ames, IA) for assistance in metabolic analyses; Bri Vidrine (Iowa State University) for providing maize *glossy2* mutant seed, and propagating this genetic stock; the Hybridoma Facility (Iowa State University, Ames, IA) for characterization of GLOSSY2 antisera.

---

**Supplemental Figure 1.** Effect of the *gl2* mutation on the extracellular cuticular lipid profiles of seedling leaves of maize at the 3-5 leaf stage. All numeric data and statistical analysis can be found in Supplemental Table 1. The data represent the average ± standard error obtained from four replicate pools, each containing five different plants.

**Supplemental Figure 2.** Effect of transgenic expression of *ZmGl2*, *ZmGl2-like* or *CER2* on the minor constituents of the extracellular cuticular lipid profiles of stems of the Arabidopsis *cer2-5* mutant plants, as compared to the non-transgenic wild-type and *cer2-5* mutant stems. Data for the C26-C32 major constituents are presented in Figure 4. All numerical data and statistical analysis are in Supplemental Table 1. The data represent mean ± standard error of 10-15 replicates.

**Supplemental Figure 3.** Extracellular cuticular lipid composition of Arabidopsis stems from the indicated genotypes. All numerical data and statistical analysis are in Supplemental Table 1. Data represent the average ± standard error (n= 10-15). The sums include alkyl chain lengths of between C12 to C32.

**Supplemental Figure 4.** Effect of the *gl2* mutation on the cellular lipidome of seedling leaves of maize at the 3-5 leaf stage. Volcano-plots of the data with the x-axis representing the log-ratios (base 2) of the relative abundance of individual metabolites as affected by the *gl2* mutation, and the y-axis represents the statistical measure of significance (p-value), evaluated by Student’s t-test; the data-points above the horizontal red dashed line are deemed statistically significant, with a p-value <0.05. The data were obtained from four replicate pools, each containing five different plants. The lipid class of each metabolite is identified by different data-symbols, and the numeral next to each data-point references the specific metabolite as identified in the numeric data presented in Supplemental Table 3. The insert image at the top-right shows an expanded view of the area encompassed by the red-dashed rectangle. The x-axis coordinate labeled as “nd” indicates metabolites whose abundance is non-detectable in one of the genotypes.

**Supplemental Figure 5.** Effect of transgenic expression of *ZmGl2, ZmGl2-like or CER2* on the stem cellular lipidome. Volcano-plots of the data, with the x-axis representing the log-ratio (base 2) of the relative abundance of individual metabolites as affected by individual transgenic events, and the y-axis represents the statistical measure of significance (p-value), evaluated by Student’s t-test. The data-points above the horizontal red dashed line are deemed statistically significantly different between the indicated genotypes, with a p-value <0.05. The insert on the top-right of each figure shows an expanded view of the area of each graph encompassed by the red-dashed rectangle. The lipid class of each metabolite is identified by different data-symbols, and the numeral next to each data-point references the specific metabolite as identified in the numeric data presented in Supplemental Table 3. The x-axis coordinate labelled “nd” indicates metabolites whose abundance is non-detectable in one of the genotypes. The transgenic events evaluated in each panel are identified in the x-axis label, and are as follows: (A) non-transgenic plants comparing *cer2-5* mutant vs wild-type; (B) transgenic expression of *CER2* in the *cer2-5* mutant background vs *cer2-5* mutant; (C) transgenic expression of *ZmGlossy2* in *cer2-5* mutant vs *cer2-5* mutant; (D) transgenic expression of *ZmGlossy2* in wild-type vs wild-type; (E) transgenic expression of *ZmGlossy2-like* in *cer2-5* mutant vs *cer2-5* mutant; and (F) transgenic expression of *ZmGlossy2-like* in wild-type vs wild-type.

## Notes

**Funding information:** This work was partially supported by the State of Iowa, through Iowa State University’s Center for Metabolic Biology, and by grants from the National Science Foundation, through awards IOS-1139489 and EEC-0813570 to BJN, and an EAPSI award OISE-1614020 to LEA.

